# Substrate stiffness engineered to replicate disease conditions influence senescence and fibrotic responses in primary lung fibroblasts

**DOI:** 10.1101/2022.09.27.509806

**Authors:** Kaj E.C. Blokland, Mehmet Nizamoglu, Habibie Habibie, Theo Borghuis, Michael Schuliga, Barbro N. Melgert, Darryl A. Knight, Corry-Anke Brandsma, Simon D. Pouwels, Janette K. Burgess

## Abstract

In idiopathic pulmonary fibrosis (IPF) there is excessive ECM deposition, increased stiffness and ultimately destruction of lung parenchyma. IPF presents mainly in the elderly, implying that senescence, a hallmark of ageing, contributes to disease progression. Several studies have reported that IPF is characterised by increased senescence and accumulating evidence suggests that structural changes, such as increased stiffness may contribute to senescence. This study therefore investigated if increased tissue stiffness could modulate markers of senescence and/or fibrosis in primary lung fibroblasts. Using hydrogels representing healthy and fibrotic stiffnesses, we cultured primary fibroblasts from non-diseased lung tissue on top of these hydrogels for up to seven days before assessing senescence and fibrosis markers. Fibroblasts cultured on stiff (±15kPa) hydrogels showed higher Yes-associated protein-1 (YAP) nuclear translocation compared to soft hydrogels. When looking at senescence-associated proteins we also found higher secretion of receptor activator of nuclear factor kappa-B ligand (RANKL) but no change in transforming growth factor-β1 (TGF-β1) or connective tissue growth factor (CTGF) expression and higher decorin protein deposition on stiff matrices. With respect to genes associated with fibrosis, fibroblasts on stiff hydrogels compared to soft had higher expression of smooth muscle alpha (α)-2 actin (*ACTA2), collagen (COL) 1A1* and *fibulin-1* (Fbln1) and higher Fbln1 protein deposition after seven days. Our results show that exposure of lung fibroblasts to fibrotic stiffness activates genes and secreted factors that are part of fibrotic responses and part of the senescence-associated secretory profile (SASP). This overlap may contribute to the creation of a feedback loop whereby fibroblasts create a perpetuating cycle reinforcing disease progression in IPF.

## 1 Introduction

The extracellular matrix (ECM) is a complex structure composed of various proteins, proteoglycans and other biological factors that are secreted and/or modified by cells embedded within this microenvironment (1). Our understanding of ECM has greatly advanced in the last decade, particularly with respect to its role in providing essential biochemical and biomechanical cues that regulate cellular functions (2, 3). During normal tissue homeostasis, fibroblasts that reside in the ECM are maintained in a quiescent state; meaning they do not proliferate but secrete, sustain, and remodel the ECM. The biomechanical cues provided by the three-dimensional ECM, that provides the microenvironment in which these cells reside, regulate cellular processes such as adhesion, migration, proliferation and differentiation (4, 5). Using specific receptors, including integrins, fibroblasts are able to sense and respond to changes in the ECM (6). The mechanical forces that are generated in response to changes in the ECM local environment generate signals within the cell that rapidly propagate and activate signalling pathways (7).

The remodelling of ECM during fibrosis leads to a change in composition, structural organisation, increased crosslinking and ultimately increased stiffness. These changes greatly influence the biochemical and biomechanical properties of the ECM, leading to alterations in cellular response such as migration and proliferation (8). Tissue damage that leads to disruption of mechanical homeostasis drives mechano-activation of quiescent fibroblasts into myofibroblasts, which is a feature of tissue wound repair (9). After successful wound repair myofibroblasts are cleared from the tissue via apoptosis to prevent excessive ECM production and contraction (10). Myofibroblasts are characterised by expression of contractile proteins such as α-smooth muscle actin (αSMA) and their ability to contract tissue. During repair they play a key role in remodelling the injured tissue and are responsible for excessive ECM deposition in fibrotic diseases such as idiopathic pulmonary fibrosis (IPF). Furthermore, as a direct effect of increased matrix stiffness, fibroblasts will produce and secrete transforming growth factor (TGF)-β1, a key fibrogenic mediator (11, 12). The increased stiffness and secretion of cytokines reinforce the activation which results in myofibroblast activation and progression of fibrosis creating a perpetuating feedback loop (13).

IPF is a fibrotic lung disease of unknown aetiology characterised by excessive ECM deposition, increased ECM stiffness and ultimately destruction of lung parenchyma (14). In IPF, the Young’s modulus (stiffness) of lung tissue has been recorded to be as high as 50 kPa, whilst in healthy lung tissue, the Young’s modulus does not exceed 5 kPa (14, 15). It is thought that IPF arises from an aberrant repair response that originates in the alveolar epithelium, and it is subsequently perpetuated by disrupted crosstalk between the epithelium and fibroblasts in lung parenchymal tissue. Recent studies have shown that senescence in both alveolar epithelial cells and fibroblasts contribute to these pathological changes in IPF (16, 17). Cells that undergo (premature) cellular senescence show marked changes in morphology, phenotype and metabolism. Most notably, irreversible cell-cycle arrest and apoptosis resistance are defining features of cellular senescence. Senescent cells remain metabolically active and acquire a dynamic pro-inflammatory secretory profile known as the Senescence-Associated Secretory Phenotype (SASP) (18). The SASP is comprised of differently secreted pro-inflammatory and pro-fibrotic cytokines which impact on neighbouring cells and the local microenvironment (19). To which degree the altered ECM in IPF contributes to disease progression is not yet fully understood, specifically in relation to cellular senescence. Evidence is emerging that increased tissue stiffness might contribute to cellular senescence. Recently it was described that stiffness could lead to increase ROS productions which is linked to cellular senescence via activation of STAT3 (20, 21). In addition, pathological stiffness leads to upregulation of TGF-β1, lysyl oxidase (LOX) and downregulation of micro-RNA-29 together with increased deposition of matricellular proteins they could potentially regulate cell-cycle arrest, apoptosis and production of cytokines through the SASP (8). Increased ECM stiffness may drive excessive ECM production creating a perpetuating feedback loop in favour of disease progression. We hypothesised that the increased stiffness of fibrotic ECM detrimentally influences fibroblast function to perpetuate disease progression in IPF. Here we aimed to investigate whether adjusting the stiffness of *in vitro* produced methacrylated-gelatine (GelMA) hydrogels resembling the stiffness of healthy and fibrotic tissues could modulate cellular senescence, the SASP or the fibrotic response in primary human lung fibroblasts.

## 2 Materials and Methods

### 2.1 Methacrylated-gelatine synthesis

Gelatine was functionalised as described before with small modifications to the original protocol (22). Ten grams medical grade 262 Bloom Type A gelatine (Gelita, Ter Apelkanaal, the Netherlands) was added to 1L 0.25M carbonate-bicarbonate (CB, Sigma-Aldrich, Zwijndrecht, the Netherlands) buffer pH 9.0 and allowed to dissolve at 50°C for 60 minutes. Heating was turned off and then 563µL methacrylic anhydride (MAA; Sigma-Aldrich, CCID: 12974) was added dropwise in three steps. Ten minutes after each step the pH was measured and corrected using a 3M NaOH (Sigma-Aldrich, CCID: 14798) solution to maintain pH 9.0. The gelatine-MAA solution was allowed to react overnight at room temperature (RT) while stirring. The next day the GelMA solution was dialysed using dialysis with molecular weight cut-off of 12 – 14kDa (Sigma-Aldrich) for up to 5 days against deionised water pH 9.0. After dialysis, GelMA solution was aliquoted in 50mL tubes, frozen with liquid nitrogen and lyophilised on a Labconco Freezone 2.5 Litre benchtop freeze-dryer (Labconco, Kansas City, MO, USA). Samples were considered dry when weights of the tubes were stable. Three batches were produced, lyophilised, re-dissolved into one large batch and then lyophilised again to minimise batch-to-batch variation.

### 2.2 Trinitrobenzenesulfonic acid assay

The degree of modification of GelMA was measured using a Trinitrobenzenesulfonic acid (TNBS) colorimetric assay (23). Two mg of dry GelMA and gelatine were dissolved in 1mL of 4% w/v sodium bicarbonate solution (Sigma-Aldrich) and 1mL of 0.5% w/v TNBS solution (Sigma-Aldrich, CCID:11045) followed by incubation at 40°C with mild shaking for two hours. Afterwards, the solution was transferred to a 7.5mL glass bottle and 3mL 6N HCL solution (Sigma-Aldrich, CCID: 313) was added to stop the reaction. Unreacted TNBS was extracted twice with diethyl ether (Sigma-Aldrich, CCID: 3283) by removing the upper phase. Absorbance (346nm) of samples was measured using a Benchmark plus UV-VIS spectrophotometer (Bio-Rad Technologies). A blank sample was generated as described above but without the addition of GelMA or gelatine and was subtracted from each measurement. The degree of functionalisation (%DoF) was calculated using the formula:

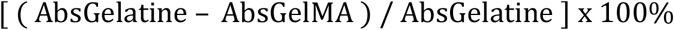

### 2.3 Fabrication of GelMA Hydrogels

GelMA samples were dissolved at 5 and 8% w/v in PBS containing 1% w/v filter sterilised lithium phenyl-2,4,6-trimethylbenzoylphosphinate (LAP; TCI Europe, Zwijndrecht, Belgium, CCID: 68384915), and kept in the dark at 37°C prior to usage. To assemble the gels, 400µL of 5 or 8% GelMA solution was added per well of a 24-well plate (Corning, New York, USA) and irradiated using a Unic nail dryer (MEANAIL© Paris, France) equipped with a UV/VIS (405nm) LED lamp for 4 minutes at 20mW/cm^2^. After irradiation, gels were allowed to swell overnight in low glucose Dulbecco’s Modified Eagle Medium (DMEM) (Lonza, Geleen, the Netherlands) supplemented with 10% foetal bovine serum (FBS) (Sigma-Aldrich), 100U/mL penicillin and 100µg/mL streptomycin (Gibco, Breda, the Netherlands), and 1% GlutaMAX (Gibco) prior to use in experiments. This media was referred to as complete growth media.

### 2.4 Mechanical testing with low load compression testing

Mechanical properties of the GelMA hydrogels were measured using Low Load Compression Testing (LLCT) as previously described (15). LLCT analyses were performed on three different locations (2.5mm apart) of each hydrogel with a 20% fixed strain rate. The stress and strain values for each measurement were determined using formulas (i) and (ii), respectively. Young’s modulus (E, i.e., stiffness) values were calculated from the slope of the linear region of the Stress-Strain graph, using formula (iii). After the peak value, the viscoelastic relaxation measurement was performed for 5 seconds, and the relaxation percentage was measured according to the formula (iv).

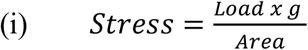

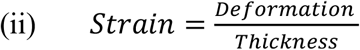

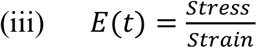

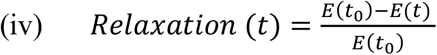

### 2.5 Human lung tissue

The study protocol was consistent with the Research Code of the University Medical Centre Groningen (https://www.umcg.nl/SiteCollectionDocuments/English/Researchcode/umcg-research-code-2018-en.pdf) and national ethical and professional guidelines (“Human tissue and medical research: code of conduct for responsible use (2011) https://www.coreon.org/wp-content/uploads/2020/04/coreon-code-of-conduct-english.pdf). Lung fibroblasts and lung tissues used in this study were derived from leftover lung material after lung surgery and transplant procedures. This material was not subject to the Medical Research Human Subjects Act in the Netherlands, and, therefore, an ethics waiver was provided by the Medical Ethical Committee of the University Medical Center Groningen (UMCG). All samples and clinical information were deidentified before experiments were performed. Fibroblasts were isolated from macroscopically and histologically normal tissue as described before (24).

Table 1 shows the characteristics of fibroblast donors used in this study: comprising 6 female and 1 male donors, 4 current smokers, 2 ex-smoker and 1 never smoker. The average age of the donors was 51.6 ± 7.87 years, all of which had normal lung capacity as demonstrated by a forced expiratory volume in 1 second/forced vital capacity (FEV1/FVC) that was above 70%.

**Table 1.** Characteristics of fibroblast donors.

| Donor # | Sex | Age | Smoking History | Pack Years | FEV1/FVC |
| --- | --- | --- | --- | --- | --- |
| 1 | F | 49 | Current | 33 | 82.3 |
| 2 | F | 50 | Never | 0 | 77.9 |
| 3 | F | 47 | Current | 30 | 73.9 |
| 4 | F | 51 | Current | 70 | 78.3 |
| 5 | F | 49 | Ex | 35 | 77.7 |
| 6 | F | 46 | Ex | 32 | 81.5 |
| 7 | M | 69 | Current | 20 | 70.0 |

F = female, M=male and Ex = former smoker

### 2.6 Cell culture

Primary LFs were cultured in complete growth media and used between passage 3 to 5 (19). LFs were routinely checked for mycoplasma infection using a PCR assay, and only used when certified negative (25). To improve attachment of fibroblasts, hydrogels were coated with a 5% w/v Bovine Serum Albumin (BSA) solution (Sigma-Aldrich) in low glucose DMEM for 24 hours prior to cell seeding. Twenty thousand LFs/gel were seeded on top of soft and stiff GelMA hydrogels in complete growth media. After 24 hours, the media was replaced with fresh complete growth media, and cells were allowed to grow for another 6 days after which supernatant, RNA, hydrogels for live/dead stain and immunofluorescent staining were harvested. Supernatants were collected, pooled, and centrifuged for 5 minutes at 300 x g to remove debris and dead cells after which the cell-free supernatants were stored at -80°C. Hydrogels in RNA lysis buffer (Machery-Nagel, Düren, Germany) were stored at -80°C until RNA isolation. Hydrogels for immunofluorescent imaging were fixed in 2% paraformaldehyde (PFA; Sigma-Aldrich) and 0.2% glutaraldehyde in PBS for 30 minutes. The hydrogels were then washed three times in PBS and stored in PBS supplemented with 1% Pen/Strep at 4°C until analyses.

### 2.7 Live and dead cell assays

Cell viability after 1 and 7 days was examined using Calcein AM (Thermo Scientific, Breda, the Netherlands, CCID: 390986) for live cells and propidium iodide (PI; Sigma-Aldrich, CCID: 104981) for dead cells. Briefly, GelMA hydrogels were washed and incubated with Hank’s Balanced Salt Solution (HBSS; Gibco) containing magnesium and calcium for 20 minutes at 37°C. A working solution was prepared containing 5 µM Calcein AM and 2 µM PI in HBSS, which was added to the GelMA hydrogels before they were incubated for 20 minutes in the dark. Images were captured using a EVOS Cell Imaging System (Thermo Scientific). ImageJ was used to create an overlay of both GFP and Texas Red channels without modifications (26). Morphological analysis was performed using CellProfiler software (Broad Institute, MIT, Cambridge MA) (27).

### 2.8 Gene expression analysis

Total RNA from GelMA hydrogels was isolated using a Nucleospin RNA isolation kit (Machery-Nagel). Media was removed and lysis buffer in a 1:3.3 sample-to-buffer ratio was added to each well before incubation for 15 minutes on ice, after which the samples (gel and lysis buffer) were stored at - 80°C. Samples were thawed on ice and mixed thoroughly before the lysis buffer was transferred to a new Eppendorf tube without the GelMA hydrogel. RNA isolation was performed according to the manufacturer’s instructions. Total RNA was quantified using the Qubit HS RNA assay kit (Thermo Scientific) according to the manufacturer’s instructions. RNA was reversed transcribed into cDNA using the ReverseAid First Strand cDNA Synthesis Kit (Thermo Scientific). DNA was amplified using GoTaq Probe qPCR Master Mix (Promega, Leiden, the Netherlands) and TaqMan Gene Expression Assay (Thermo Scientific) on a ViiA7 Real-Time PCR System (Applied Biosystems, Breda, the Netherlands) with the relevant predeveloped primers listed in table 2 (Thermo Scientific).

**Table 2.** Primers and probes with designated exon boundaries

| Gene | Number | Exon boundary |
| --- | --- | --- |
| 18S | Hs99999901_s1 | 1-1 |
| CDKN2A | Hs00923894_m1 | 2-3 |
| CDKN1A | Hs00355782_m1 | 2-3 |
| P53 | Hs01034249_m1 | 10-11 |
| IL-6 | Hs00174131_m1 | 4-5 |
| CXCL8 | Hs00174103_m1 | 1-2 |
| DCN | Hs00370385_m1 | 7-8 |
| ACTA2 | Hs00426835_g1 | 2-3 |
| COL1A1 | Hs00164004_m1 | 1-2 |
| FN1 | Hs01549976_m1 | 8-9 |
| FBLN1 | Hs00242545_m1 | 13-14 |
| FBLN1C | Hs00242546_m1 | 14-15 |
| LOX | Hs00942483_m1 | 5-6 |
| TGF-β1 | Hs00998133_m1 | 6-7 |
| CTGF | Hs00170014_m1 | 4-5 |

18S ribosomal RNA = 18S, Cyclin-dependent kinase inhibitor 2A = CDKN2A, Interleukin 6 = IL-6, Chemokine ligand 8 = CXCL8, Decorin = DCN, Actin alpha 2 = ACTA2, Collagen 1α1 = COL1A1, Fibronectin = FN1, Fibulin-1 = FBLN1, Lysyl oxidase = LOX, Transforming growth factor β1 = TGF-β1, Connective tissue growth factor = CTGF

### 2.9 Immunofluorescence

For immunofluorescence cells were washed twice with PBS and fixed with 2% PFA/0.2% glutaraldehyde (Sigma-Aldrich) in PBS for 20 minutes at room temperature. Fixed cells were permeabilized with 0.5% Triton X-100 in PBS for 10 minutes and incubated with 2.2% bovine serum albumin (BSA) for 30 minutes. Next cells were incubated with primary antibodies for 16 hours: rabbit polyclonal to YAP (GeneTex Cat# GTX129151, 0.3 µg/ml, RRID:AB_2885910), mouse monoclonal to P16 (Abcam Cat# ab54210, 2.5µg/ml, RRID:AB_881819), rabbit polyclonal to Decorin (Proteintech Cat# 14667-1-AP, 0.9 µg/ml, RRID:AB_2090265) 14667-1-AP; ThermoFisher), rabbit polyclonal to fibulin-1 (Thermo Fisher Scientific Cat# PA5-103841, 3.33 µg/ml, RRID:AB_2853174) in PBS + 0.1% Triton X-100 and 2.2% BSA. After three washes with PBS containing 0.5% Tween-20 samples were incubated with fluorescent labelled secondary antibodies containing 4′,6-diamidino-2-phenylindole (DAPI; Roche Cat# 10236276001, 1.0 µg/ml) for 30 min. Actin was visualized by incubating with TRITC labelled-Phalloidin (Sigma-Aldrich Cat# P1951, 100nM, RRID:AB_2315148) in PBS for 30 min. Slides were mounted in Citifluor (Agar Scientific, Stansted, UK). Fluorescent photomicrographs were acquired using a SP8 scanning confocal microscope equipped with photomultiplier tubes (PMTs) (Leica Microsystems, Amsterdam, the Netherlands) and a HC PL APO CS2 63x/1.4 oil immersion objective at room temperature. Fluorescent images were loaded in ImageJ and subsequently split into the separate channels. The DAPI channel was used to count the number of nuclei and determine the nucleic area. This was done using the particle Analyzer with particles size set at 50 – Infinity. Then, a mask was made of the nuclear area. In the YAP channel total positive area was measured and by using the nuclear mask also the nuclear area was determined. The nuclear fraction was calculated as follows: total nuclear YAP was divided by total YAP to calculate the ratio between total and nuclear YAP. Matrix ECM patterns were analysed using The Workflow Of Matrix BioLogy Informatics (TWOMBLI) as described before (28).

### 2.10 Enzyme linked immunosorbent assay

Secreted IL-6 (DY205), CXCL8 (DY208), DCN (DY143), TGF-β1 (DY240), Osteoprotegerin (OPG; DY805), receptor activator of nuclear factor kappa-B ligand (RANKL; DY626) and Connective Tissue Growth Factor (CTGF) (DY9190-05) were measured using Human DuoSet Enzyme linked immunosorbent assay (ELISA) kits (R&D Systems, Abingdon, UK) according to the manufacturer’s instructions. Total RANKL was calculated by measuring free RANKL and in complex with OPG using an in house developed OPG/RANKL complex sandwich ELISA by using a human capture RANKL antibody (part of cat#DY626, R&D Systems) and a human detection OPG antibody (part of cat#DY805, R&D Systems) (29). Total OPG was calculated by measuring free OPG and bound OPG. To normalise ELISA data, cell numbers from ten fields of DAPI positive cells per donor on soft (±5kPa) and stiff (±15kPa) GelMA hydrogels were counted, and the total number of cells calculated for the overall GelMA hydrogel surface (1.9cm^2^) area.

### 2.11 Statistical Analyses

Statistical analyses were performed using IBM SPSS Statistics for windows version 25 (IBM Corp., Armonk, N.Y., USA) and evaluated using Wilcoxon matched pair signed rank test for comparison between soft GelMA hydrogels and stiff GelMA hydrogels. Data were considered statistically significant at p < 0.05 unless stated otherwise. D’Agostino test was used to examine normality of the data and when necessary, log transformation applied. Correlations were assessed by calculating a Pearson correlation coefficient.

## 3 Results

### 3.1 Mechanical properties of GelMA hydrogels

Three different GelMA batches were produced and the methacrylation consistency was tested to determine if they could be combined into one large batch. The three different GelMA batches prepared for this study had a methacrylation efficiency of 94.58 ± 1.98%, 97.84 ± 1.14% and 89.77 ± 2.04% respectively (figure 1A). The combined batch had a methacrylation efficiency of 88.22 ± 1.32%. Next, the Young’s modulus was measured across a range of different percentage GelMA hydrogels of up to 10% GelMA (data not shown) to select the hydrogel composition with the stiffness that best mimicked healthy (±5kPa) and fibrotic (±15kPa) lung tissue. A hydrogel made of 5% wt/v GelMA resembled healthy tissue with Young’s modulus of 3.94kPa ± 0.87kPa whereas 8% wt% hydrogels most resembled stiff tissue with a Young’s modulus of 11.34kPa ± 3.82kPa (figure 1B). To further characterise these hydrogels, relaxation was measured to see if the percentage of GelMA has an impact on relaxation. After initial compression of 20%, the dissipation of force was followed for a duration of 5 seconds (figure 1C). The relaxation values were 4.54 ±1.60% for 5% (soft) and 4.00 ± 0.32% for 8% (stiff) GelMA hydrogels.

**Figure 1.**
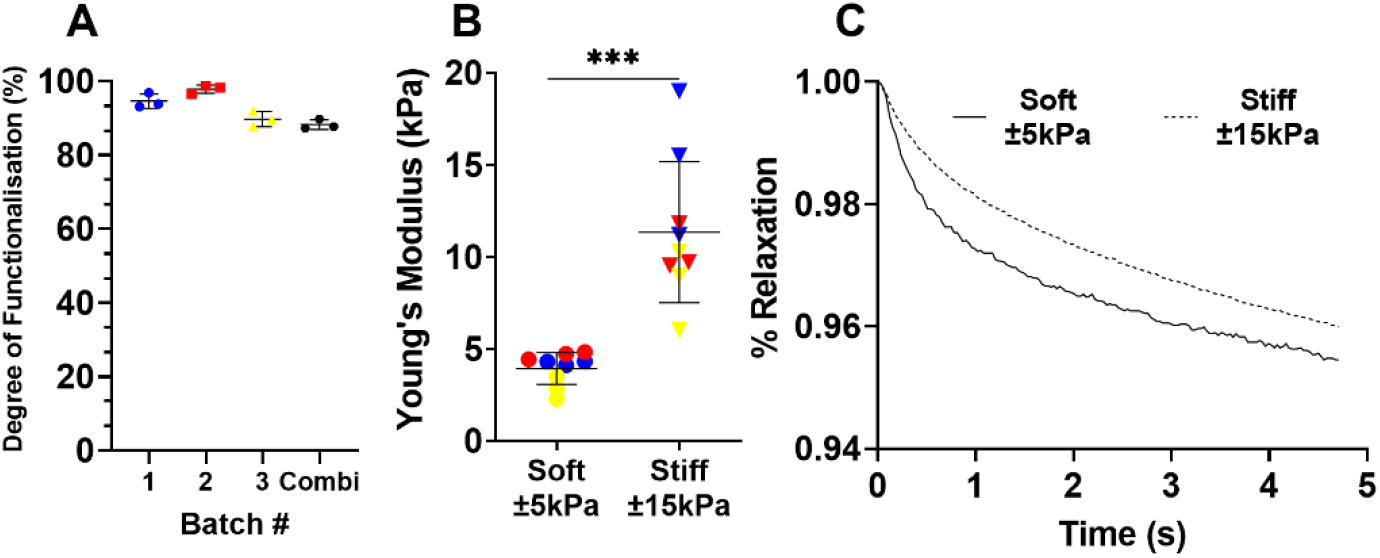
The methacrylation efficiency was measured to demonstrate minimal variety before being combined into one large batch. Each dot represents a technical replicate of a batch. Data expressed as mean ±SD. Rheological properties were characterised by the Young’s modulus of 3 different series (B; blue, red and yellow) produced from three GelMA hydrogel solutions used in this study and the relaxation (C) over a period of 5 seconds. N=3 with each dot representing a technical replicate of a unique sample. Data is expressed as mean ± SD in panel A-B, and mean in panel C (n=3). Data were considered significant at p < 0.05

### 3.2 Morphological differences of fibroblasts on soft and stiff GelMA hydrogels

To characterise responses of fibroblasts to their environment, fibroblasts were cultured on soft (±5kPa) and stiff (±15kPa) hydrogels and their morphology was observed after 1 and 7 days of culture. Fibroblasts seeded on both soft and stiff hydrogels exhibited a similar morphology after 24 hours of culture (figure 2A-B). After 7 days of culture, fibroblast morphology was also similar on both soft and stiff hydrogels (figure 2A-B day 7). After observing a round morphology in some donors after 24 hours a live/dead stain was performed to rule out apoptosis, which revealed live cells with no dead cells evident after 1 or 7 days of culture on either hydrogel (figure 2C-D). Total fibroblast cell number after 7 days of culture was higher on stiffer hydrogels compared to soft hydrogels (figure 2E) which is in concordance with literature in response to a stiff matrix. Morphology analysis using CellProfiler did not demonstrate major differences between fibroblast on day 1 or 7 and between soft and stiff hydrogels (supplementary figure S1).

**Figure 2.**
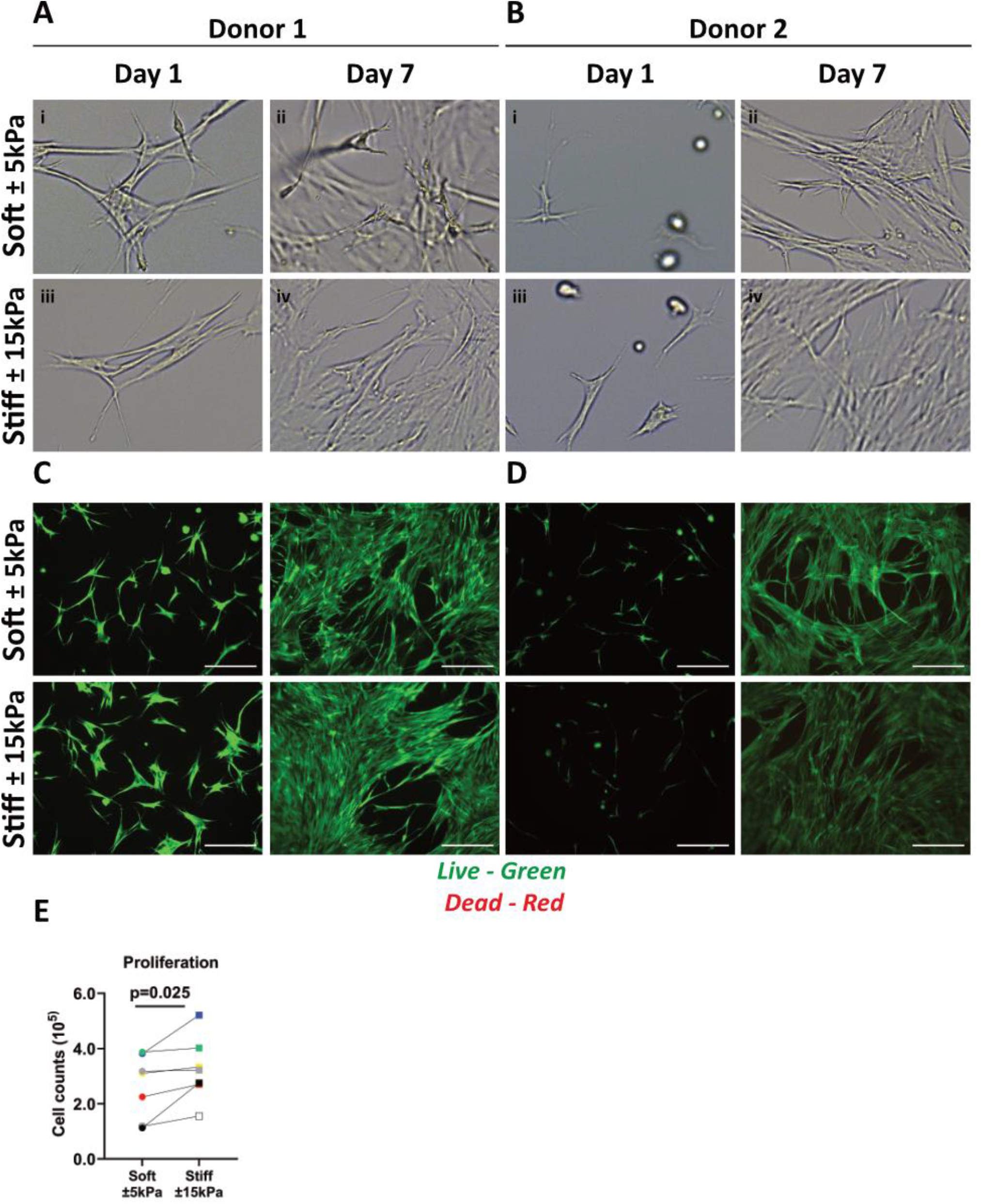
Primary human lung fibroblast morphology, live/dead staining and proliferation on soft (3.94 ± 0.87 kPa) and stiff (11.34 ± 3.82 kPa) GelMA hydrogels. A-B) Representative brightfield microscopy images of cells 24 h post seeding and after 7 days of culture on top of soft and stiff hydrogels (digital magnification). Letters i-iv show areas of interest with different morphologies between donors. C-D) Live (green) and dead (red) stain of two donors at day 1 and 7 of culture. E) Total cell numbers on soft (3.94 ± 0.87 kPa) and stiff (11.34 ± 3.82 kPa) hydrogels after 7 days of cell culture. Images are representative of responses seen in 7 unique donors. (n=7) Scale bar = 250 µm. Each donor has a different unique colour (blue, red, yellow, green, black, white, and grey).

### 3.3 Increased total YAP content and organised F-actin formation in fibroblasts on stiffer matrices

Next, the mechanosensory response of fibroblasts to their environment was examined using a dual stain for Yes-associated protein 1 (YAP) nuclear translocation and F-actin fibre formation. The YAP nuclear fraction was higher on stiff hydrogels compared to soft hydrogels (figure 3A). Figure 3B shows the nuclear translocation of YAP on soft and stiff hydrogels on day 7. F-actin cytoskeleton arrangement in fibroblasts on soft hydrogels was composed of both diffuse and organised F-actin fibres, while fibroblasts on stiffer hydrogels display bundled organised F-actin fibres (figure 3B), indicative of their responsiveness to the substrate stiffness. To quantify the differences in ECM patterns between soft and stiff hydrogels we measured several parameters using the TWOMBLI plugin. Figure 3C demonstrates that there is a clear increase in total area of F-actin fibres in fibroblasts cultured on stiff hydrogels. This is supported by the reduction in space between the fibres (Figure 3D) as well as the increase in F-actin fibre length (Figure 3E). Lasty, we measured a strong increase in the high-density fibre network, which confirms areas with increased tight f-actin bundles formed in response to stiffness (figure 3F).

**Figure 3.**
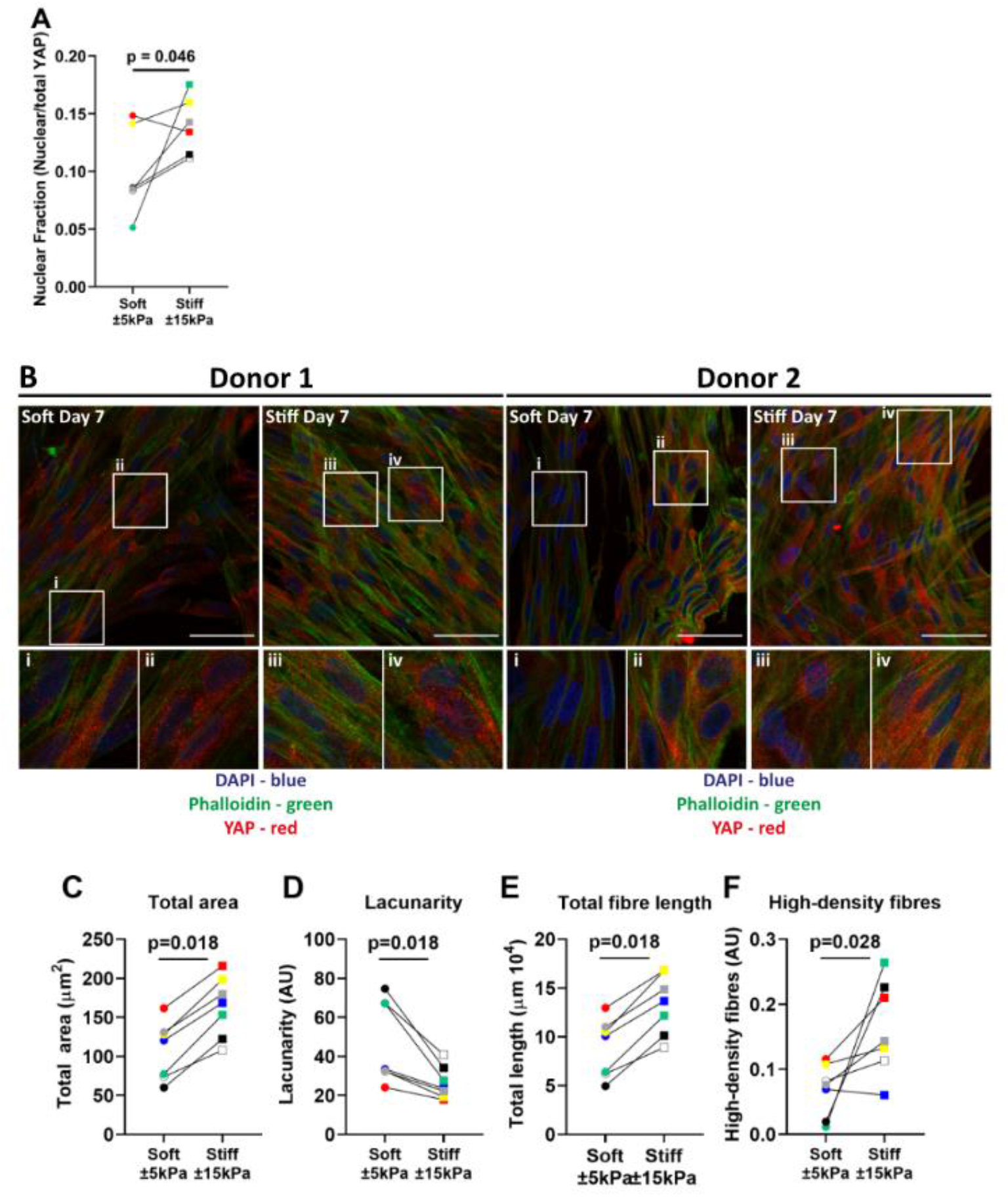
YAP translocation and F-actin stress fibre formation in primary human lung fibroblasts on soft (3.94 ± 0.87 kPa) and stiff (11.34 ± 3.82 kPa) hydrogels. Hydrogels after 7 days of culture were fixed and fibroblasts were stained for YAP and F-actin. A) Quantification of YAP nuclear fraction B) representative fluorescent images of DAPI (Blue), YAP (Red) and F-actin (green). i-iv) White boxes show the region of interest that is magnified. TWOMBLI analyses of phalloidin stained fibroblasts showing total area (C), lacunarity (D), fibre length (E) and high-density fibre networks (F) (calculated using the high-density matrix algorithm). Images are representative of responses seen in 7 unique donors. (n=6-7) Scale bar = 50µm. Each donor has a different unique colour (blue, red, yellow, green, black, white, and grey)

### 3.4 Higher matrix stiffness does not contribute to cellular senescence in fibroblasts

After confirming that fibroblasts respond to their environment, we explored our hypothesis that increased stiffness may contribute to the senescent state. Therefore, several biomarkers for senescence were analysed in fibroblasts grown on soft and stiff hydrogels mimicking healthy and diseased tissue stiffnesses. Gene expression analysis of early-stage marker *p21*^*Waf1/Cip1*^ did not demonstrate a difference while late-stage senescence marker *p16*^*Ink4a*^ demonstrated a trend (p=0.075) towards an increase between soft and stiff hydrogels (figure 4C-B). Interestingly, *p53* demonstrated a trend towards a decrease in fibroblasts on stiff hydrogels (Figure 4C). To further characterise cellular senescence, both gene expression and secretion of the well characterised SASP factors IL-6 and CXCL8 were measured (figure 4D-G). Gene expression of *IL-6* was significantly higher on stiffer hydrogels (figure 4D). However, the secretion of IL-6 did not change between soft and stiff hydrogels. Both CXCL8 gene and protein expression did not exemplify a difference between soft and stiff hydrogels after 7 days of culture (figure 4F-G).

**Figure 4.**
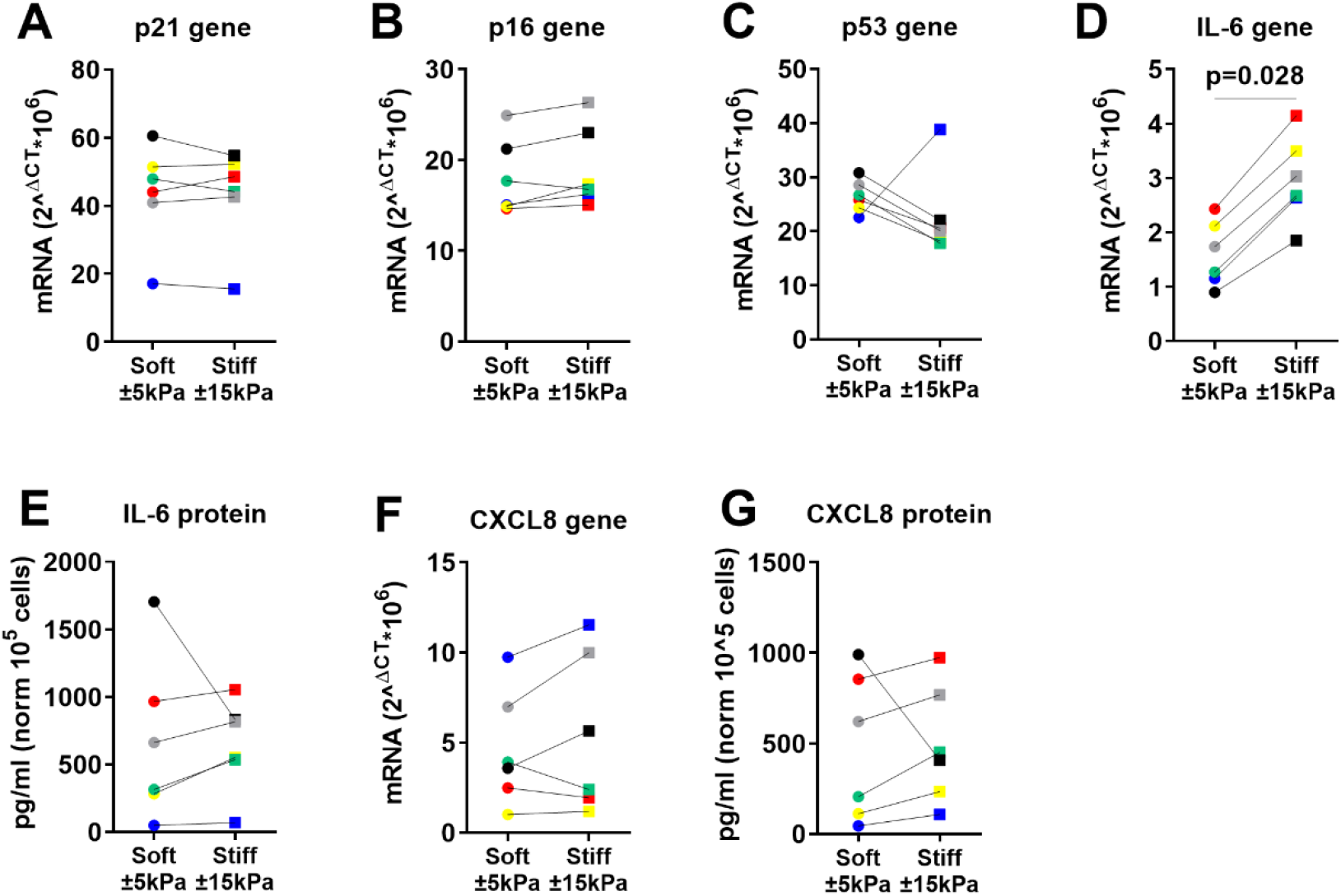
Primary human lung fibroblast markers of cellular senescence in response to surface stiffness. A-C) Gene expression of the cell-cycle inhibitors *p21*^*Waf1/CIp1*^ *and p16*^*Ink4a*^, *p53* and known SASP factors *IL-6* and *CXCL8* after 7 days of culture, data were normalised against 18S and expressed as 2^-ΔCT^ × 10^6^. D-G) Levels of SASP cytokine production in supernatant, data were normalised to cell numbers and are expressed as pg/mL per 10^5^ cells. Differences between soft (3.94 ± 0.87 kPa) and stiff (11.34 ± 3.82 kPa) hydrogels were analysed by a Wilcoxon matched pairs signed rank test (n = 6) and p<0.05 was considered significant. Each donor has a different unique colour (blue, red, yellow, green, black, and grey).

### 3.5 Higher matrix stiffness induces a secretory phenotype in primary fibroblasts

Our data indicated that there was no greater activation of the cell-cycle inhibitors *p16*^*Ink4a*^ and *p21*^*Waf1/Cip1*^ on stiffer hydrogels compared to softer hydrogels, but rather an increase in proliferation. To further characterise the activation of a secretory phenotype we measured several factors that have been reported to be part of the senescent phenotype and/or to be part of a fibrotic response (18, 30-32). Gene expression and secreted protein levels of the proteoglycan DCN were higher on stiff compared to soft hydrogels (figure 5A). Next, RANKL was measured as it was recently shown to be secreted by senescent fibroblasts in COPD (33). RANKL secretion was significantly higher at day 7 on stiffer hydrogels while its decoy receptor OPG showed a trend towards an increase at day 7 on stiff hydrogels (figure 5C-D). Additionally, TGF-β1 gene expression and protein secretion were measured. TGF-β1 gene expression demonstrated a significant decrease on stiff hydrogels compared to soft hydrogels (figure 5E-F) while there was no change in protein secretion (figure 5G). Finally, CTGF gene expression and secretion were measured, which demonstrated no difference for secretion between soft and stiff hydrogels at day 7 while gene expression demonstrated a trend towards a decrease (figure 5G-H). To explore if the OPG-RANKL axis serves as a mechanism that potentially could have an effect on both fibrosis and senescence, we explored the correlation between OPG-RANKL and other secreted factors (Figure 5I). Correlations for both OPG and RANKL were observed with IL-6, CXCL8 and TGF-β1. CTGF was correlated with OPG but not RANKL, and no correlations were observed with DCN. Additionally, we stained the cells on the surfaces of the hydrogels for DCN protein levels. Figure 5J shows the DCN stain on both soft and stiff hydrogels after 7 days in culture. DCN displayed a strong visual increase on the stiffer hydrogels in comparison to the soft hydrogels.

**Figure 5.**
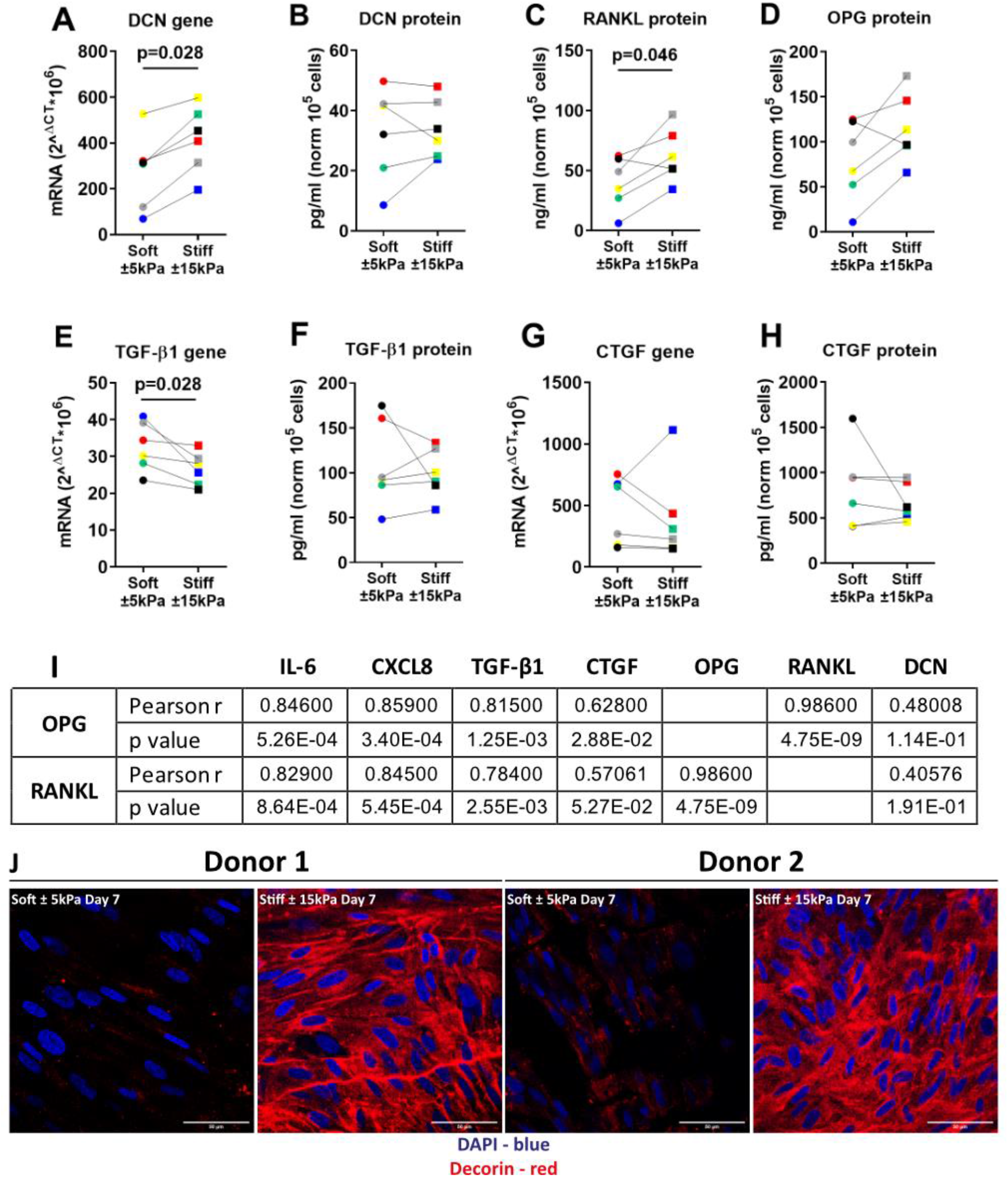
Stiffness induced a secretory phenotype in Primary human lung fibroblasts cultured on stiff matrices. A-B) DCN gene expression and secreted protein were measured after 7 days of culture on soft (3.94 ± 0.87 kPa) and stiff (11.34 ± 3.82 kPa) hydrogels. C-H) Levels of secreted factors RANKL, OPG, CTGF and TGF-β1 including TGF-β1 and CTGF gene expression after 7 days of culture on soft and stiff hydrogels. I) Correlation was calculated between OPG, RANKL and all other secreted factors. Panel J shows a fluorescent staining for DCN, DAPI was used for identification of cell nuclei. Illustrated photographs are representative images of in total 7 unique donors. Gene expression data were normalised against 18S and expressed as 2^-ΔCT^ × 10^6^. Secreted matrix proteins and cytokines were normalised to cell counts and expressed as pg/mL or ng/ml 10^5^ cells. Differences between soft and stiff hydrogels were analysed by a Wilcoxon matched pairs signed rank test. Correlation was calculated using Pearson correlation coefficient. (n=6) Scale bar = 50µm. Each donor has a different unique colour (blue, red, yellow, green, black, and grey).

### 3.6 Matrix stiffness activates a profibrotic response in primary fibroblasts

To further characterise the fibrotic response of fibroblasts on stiff hydrogels we measured several fibrosis-associated genes. Previous studies have reported that higher expression of these genes is associated with persistent fibroblast activation and higher ECM deposition in fibrosis. Lung fibroblasts on stiff hydrogels exhibited higher α-SMA and COL1α1 expression after 7 days of culture compared to soft hydrogels (figure 6A-B). The expression of FN1 was significantly lower after 7 days on stiff hydrogels (figure 6C). Interestingly, LOX gene expression was lower in cells grown on stiff hydrogels compared to soft hydrogels (figure 6D). Fibulin-1 gene expression was higher after 7 days on stiff hydrogels (figure 6E). However, FBLN1C gene expression was not different between cells grown on soft or stiff hydrogels (figure 6F). The final characterisation was an immunofluorescent staining for deposited FBLN1 on both soft and stiff hydrogels at day 7. We visualised a higher deposition of FBLN1 on stiff hydrogels in comparison to soft hydrogels (figure 6G).

**Figure 6.**
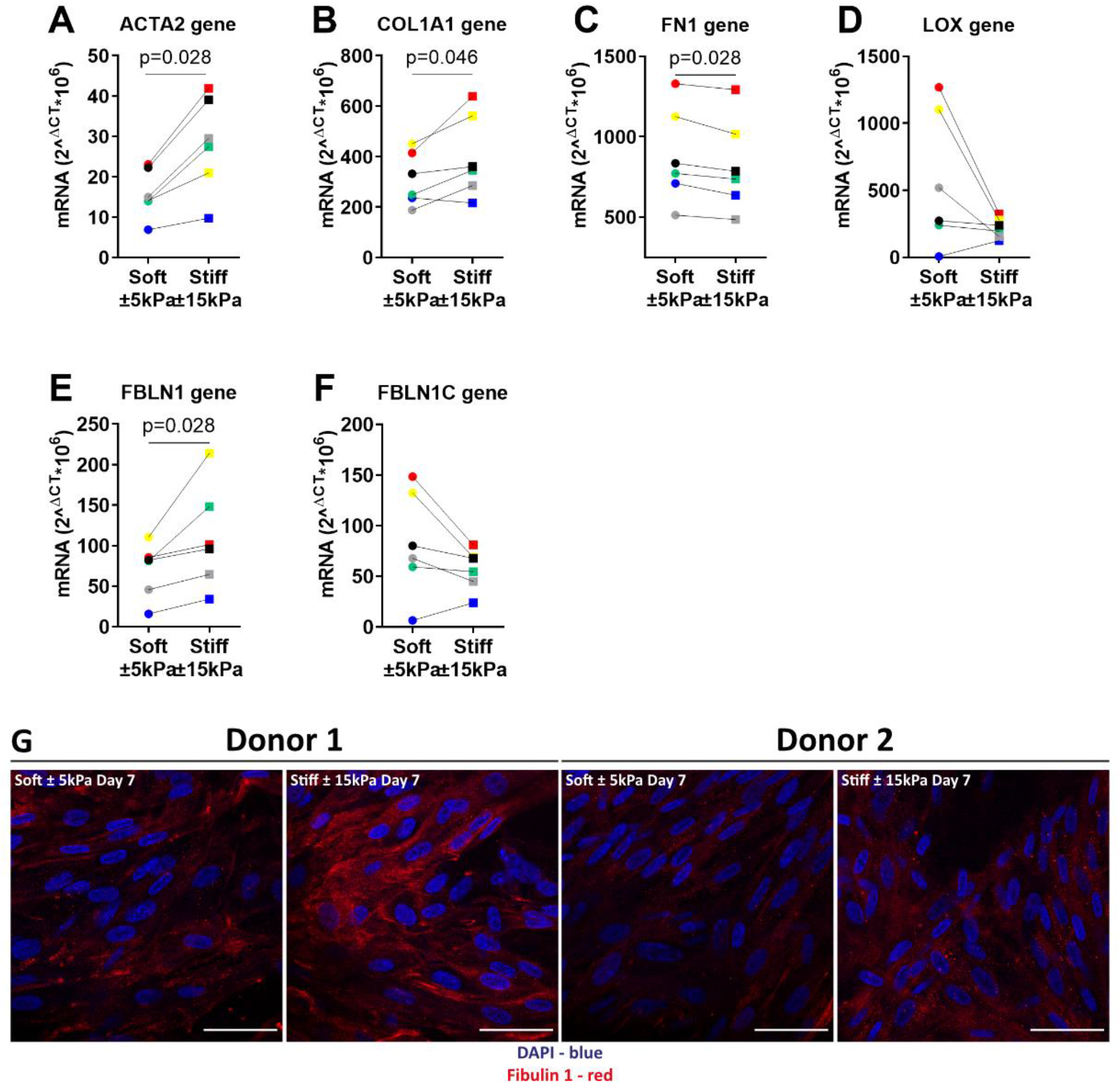
Primary human lung fibroblasts exhibit an activated and profibrotic phenotype on stiff (11.34 ± 3.82 kPa) compared to soft (3.94 ± 0.87 kPa) hydrogels. To characterise the fibrotic response of fibroblasts on hydrogels, gene expression levels for fibroblast activation α-SMA and matrix proteins COL1α1, FN1, LOX, FBLN1 and FBLN1C were measured at day 7 (A-F). Finally, deposited FBLN1 (Red) was stained using fluorescent labelled antibodies and DAPI for identification of cell nuclei in panel G. Illustrated photographs are representative images of in total 7 donors. Differences between soft and stiff hydrogels were analysed by a Wilcoxon matched pairs signed rank test (n = 6). Scale = 50 µm. Each donor has a different unique colour (blue, red, yellow, green, black, and grey).

## 4 Discussion

During fibrotic disease, the ECM undergoes tremendous change that influence the composition and stiffness of the tissue. These changes have a substantial effect on cell function, and it has been reported that fibrotic ECM is able to create a profibrotic positive feedback loop (34). In the present study we created hydrogels reflecting the pathological stiffness measured in fibrotic diseases, such as IPF, and examined the impact on cellular responses of lung fibroblasts focusing on markers of senescence, including the SASP and fibrosis. Pathological stiffness did not affect *p16*^*Ink4a*^,*p21*^*Waf1/Cip1*^ or *p53*, the main markers of senescence, instead it resulted in higher expression of genes/proteins associated with the SASP and fibrosis such as *IL-6, ACTA2, COL1A1, FBLN1* and *DCN* compared to soft matrix. However, some markers including IL-6, OPG & RANKL are identified to be both expressed/secreted during cellular senescence and different stages of fibrosis. Collectively, the data presented in this study suggest that exposure of fibroblasts to a surface of pathological stiffness modulates gene expression and secreted factors that are both involved in cellular senescence and the fibrotic response in fibroblasts.

Methacrylated-gelatine gels are well characterised hydrogels that support growth of a large variety of cells, that are biodegradable, that allow cell adhesions through Arg-Gly-Asp (RGD) sequences and allow for precise tuning of mechanical properties. Precise tuning of the degree of functionalisation, gelatine concentration and number of crosslinks results in high reproducibility and controllability of biophysiochemical properties (35-37). These characteristics made these hydrogels ideal for our study. In this study we generated two hydrogels with a different Young’s modulus while the initial stress relaxation behaviour did not significantly change. This was not unexpected as both hydrogels were almost equal in their constituent biomaterials, reflecting the difference in percentage of GelMA used. This observation authenticates that the responses of the fibroblasts measured herein could be contributed to the differences in Young’s modulus only. However, more focus is directed towards the effect of stress relaxation and how it may influence cell behaviour (38, 39). Tests on tissues such as liver, skin, muscle, and breast reveal that these organs dissipate stress ranging from milliseconds to hundreds of seconds (40, 41). Changes in viscoelastic properties have been associated with disease progression including cancer. Substrate stiffness has a robust effect on cell function and regulates cell viability, proliferation, migration, differentiation, and cell-matrix adhesion (4). Higher proliferation, contractility and activation of fibroblasts are all reported as responses to a stiffer environment (42-44). In this regard, the proliferation and high viability of fibroblasts grown on both soft and stiff hydrogels observed in this study were comparable to observations from previous investigations (43, 45, 46). Furthermore, the morphology of fibroblasts on soft gels was comparable to fibroblasts on stiff gels. This is in contrast to some literature where it is shown that there is little spread of cells on soft stiffness (45). However, the stiffness representing a soft matrix is often much lower than used in this study and is more in the range of 0.5kPa. in addition, culture times were also different which potentially contributes to the morphology as fibroblasts produce matrix of their own which facilitates attachment. Furthermore, differences in gel composition may also contribute to this effect, for example polyacrylamide has no RGD sequences as are present in GelMA due to its biological origin, and this may impact on the ability of the fibroblasts to attach and spread on the gel surface.

In response to mechanical cues such as stiffness, fibroblasts organise their actin fibres and translocate YAP into the nucleus leading to cellular activation and transcription of genes in many cell types (47-49). The data in this study showed that total YAP content in fibroblasts is higher on stiff compared to soft hydrogels and that organised bundled F-actin fibres are formed, which is indicative of the cells responding to the mechanical environment. However, there was an apparent lack of nuclear staining for YAP at day 7 in the fibroblasts on either hydrogel, particularly in regions of higher cell density. This may be explained by the fact that YAP becomes less mechanosensitive and remains in the cytoplasm as a response to cell-cell interactions (50, 51). Furthermore, while increased nuclear translocation is well documented in response to a stiff environment, the increase in total YAP is not yet clarified. In fact, stimulation with TGF-β1 demonstrated a reduction in YAP mRNA expression in the first 24 hours while protein levels did not change (52). Moreover, the formation of organised bundled actin may also contribute to this effect by altering the shape of the nuclear envelope which potentially changes the environment for transcription and thus impacting on fibroblast function (53). Little is known about the interplay between an altered ECM and cellular senescence, therefore after confirming that, in our model, fibroblast respond to their substrate, we investigated what the contribution was of increased stiffness on cellular senescence and the SASP. The results from the present study provided evidence that there was an increase of proliferation and modulation of some markers of senescence in response to pathological stiffness at day 7. Higher cell proliferation and migration, as reflected in our study, has been widely documented as a result of increased matrix stiffness and is part of an important response to injury in wound repair (54, 55). Interestingly, we also observed an increase in *IL-6* gene expression and a trend towards an increase in CXCL8 protein secretion in the stiffer environment. Both pro-inflammatory cytokines are involved in wound repair and are identified as part of the SASP, which plays an important role in fibrosis (18, 56). Furthermore, there is evidence suggesting that the SASP can be activated independently of cell-cycle inhibitors *p16*^*Ink4a*^ and *p21*^*Waf1/Cip1*^ in cellular senescence (57, 58). Activation of the SASP might be an early indicator of senescence and precedes the irreversible cell-cycle arrest of premature cellular senescence.

Activation of a secretory phenotype is beneficial for tissue repair, but it can also be detrimental and contribute to disease progression in fibrosis (59). Our results showed activation of a secretory phenotype in response to increased surface stiffness, including a strong correlation of RANKL and OPG with other secreted factors, but not DCN. All of these factors have been recognised to be part of the SASP and/or to be part of the fibrotic response. DCN is recognised for its role in many different biological processes and recently DCN has been found negatively correlated with *p16*^*Ink4a*^ in senescence in COPD (60). Our results suggest, in the context of fibrosis, that DCN deposition is in response to stiffer hydrogels, further supporting the important role DCN plays in fibrosis (32, 61). CTGF is an important mediator in tissue remodelling and has been associated with senescence, fibrosis and cancer (62, 63). CTGF can be regulated via its main inducer TGF-β1 or can be negatively regulated by DCN and OPG (64, 65). Given the fact that, in our study, TGF-β1 secretion was unaltered and an increase in DCN secretion was observed, it was not surprising that no change in CTGF secretion was found in response to stiffness. CTGF has also been reported to be downstream of YAP, and we reported there was more YAP retention in the cytoplasm potentially contributing to less upregulation in response to the stiff environment (66, 67). In addition, there was downregulation of CTGF gene expression in response to stiff hydrogels confirming the findings that protein secretion was unaltered. The RANKL-OPG system is essential for bone metabolism, where OPG acts as a decoy receptor for RANKL, regulating development and activity of osteoclasts. OPG is currently used as a biomarker for liver fibrosis (18, 68), while RANKL was recently identified to be part of the SASP in senescent COPD fibroblasts (33). In IPF the exact role of OPG is not fully understood but evidence suggests it may play a role in active fibrosis and associates with disease progression (69, 70). Our correlation analysis between RANKL, OPG and other secreted factors demonstrated a strong correlation. These factors, such as TGF-β1 and CTGF, are part of both the SASP and fibrotic response. It might be possible that RANKL as a SASP and OPG as a fibrotic-associated marker create a regulatory pathway between the senescent and fibrotic response in stiff lung tissues. Moreover, CTGF has been identified to bind to OPG and might be a potential fourth factor in the RANK-RANKL-OPG system, potentially contributing to the regulatory mechanism as well (63, 65, 71, 72). This mechanism might also contribute to the fact that we did not measure any difference in protein secretion, despite lower mRNA levels. Taken together these data suggest that the RANKL-OPG-CTGF system may contribute to secreted factor regulation, increased ECM deposition and potentially contribute to the progression/perpetuation of fibrosis in response to a stiff tissue environment as central regulator.

Fibroblasts can be activated in response to increased stiffness leading to higher expression of α-SMA and matrix proteins, which is part of a normal wound repair response after injury. Our results indicate that that in response to stiffer hydrogels fibroblasts show upregulation of several fibrosis associated genes including *ACTA2* and *COL1A1*. Increase in *FBLN1* expression and deposition on stiffer hydrogels was expected and is in line with what has been reported in literature (73, 74). However, the lack of response of *FBLN1C* gene expression was unexpected as *FBLN1C* has been reported to be implicated in fibrosis and lung injury (75, 76). However, it has been reported that TGF-β1, which is unaltered in our study, downregulates *FBLN1* gene expression and promotes FBLN1 deposition in airway smooth muscle cells (77). This could indicate that FBLN1/C regulation might be cell-type and disease specific. Interestingly, FN1 which is associated with FBLN1 was decreased on stiffer hydrogels after 7 days. This was in contrast with literature where FN1 was shown to be an important mediator of early wound repair, collagen fibril formation and fibrosis (78). Additionally, Zhou and colleagues demonstrated that FN1 expression increased when primary lung fibroblasts were cultured for up to 48 h on stiff matrixes (79). However, it is possible that FN1 peaked early in the time-course of our experiment, especially given its role in the initial phase of wound repair and had subsequently decreased at the time we measured it. Finally, *LOX* gene expression demonstrated a fine regulation by stiff hydrogels regardless of gene expression on soft hydrogels. This effect was unexpected as it was reported that there is no change in gene expression in non-IPF versus IPF. However, on protein level it was reported that LOX deposition was decreased in IPF tissue (80). Therefore, it would be appropriate to determine enzyme activity as gene expression and protein expression do not always correlate.The experimental strategy used in this study was designed to measure the impact of stiffness on the senescent phenotype in fibroblasts after 7 days in culture. While there was modulation of some markers of cellular senescence, other studies investigating senescence induce this by stimulating the cells with paraquat or hydrogen peroxide (20, 60). It is possible that the early senescence marker *p21*^*Waf1/Cip1*^ reached its highest expression before we assessed the expression at day 7, while still being too early to detect high levels of *p16*^*Ink4a*^.In several of our datasets there appear to be data points driving the lack of statistical difference, however, these are not attributed to a single donor but rather represent different donors in each dataset. This variability can be attributed to the biological heterogeneity of the primary lung fibroblasts used in this study. Cell numbers at the start of the study were optimised for measuring senescence readouts. This made the assessment of YAP translocation challenging as the density of fibroblasts on day 7 were higher than optimal for this readout. In addition, YAP translocation is a swift process as cytoplasmic/nuclear levels can be stable within 1 day after changing the stiffness (81). This could have implications for measuring YAP translocation as the regulatory effect may have been partially diminished demonstrating a weaker effect.

In conclusion, we have established that the culture of human lung fibroblasts on matrices that mimic pathological stiffness of IPF tissues for up to seven days leads to activation of genes and secretion of matrix proteins and cytokines that are part of the fibrotic response, with overlap of SASP factors (figure 7). Our data suggest that there may be an overlap between cellular senescence/SASP and the fibrotic response, indicating that the fibroblasts may acquire a pre-senescent and / or activated phenotype. The modulation of cell function in response to pathological stiffness may ultimately lead to the creation of a negative feedback loop in which pre-senescent and/or activated fibroblasts reinforce each other causing a perpetuating cycle driving aberrant ECM accumulation and disease progression in IPF.

**Figure 7.**
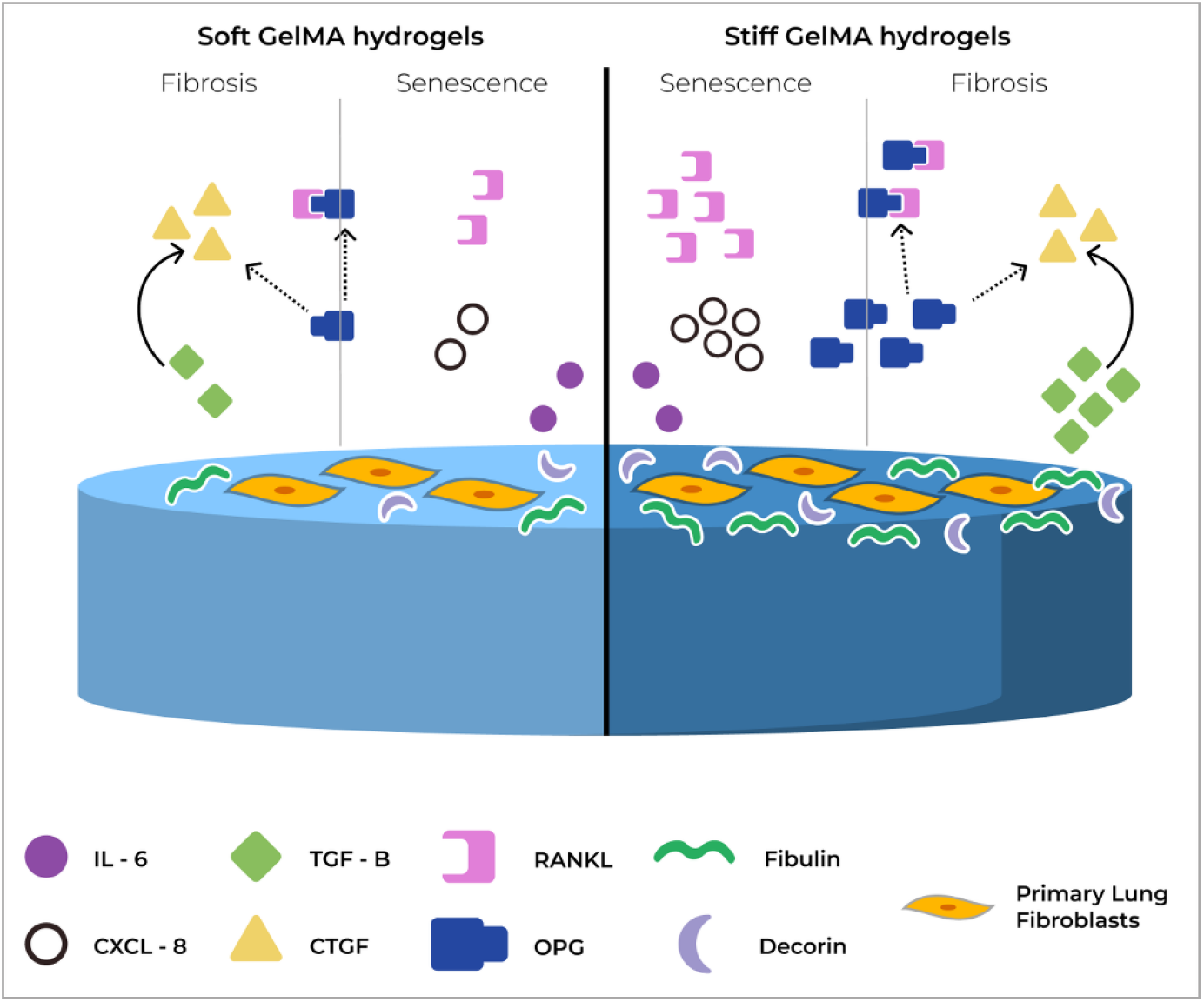
Summary of proposed mechanism for OPG in senescence and fibrosis. In response to stiffness, Primary human lung fibroblasts show increased proliferation, higher deposition of ECM proteins and higher secretion of SASP and fibrosis related cytokines. OPG is proposed to act as a regulatory mechanism between senescence and fibrosis by binding to RANKL and CTGF, and potentially inhibiting their function.

## 6 Author Contributions

K.E.C.B., M.N., M.S., D.A.K., C.A.B., S.D.P and J.K.B conceived and designed research; K.E.C.B and M.N performed experiments; K.E.C.B., M.N., H.H and T.B analysed data; K.E.C.B., M.N., H.H., M.S., B.N.M., D.A.K., C.A.B., S.D.P and J.K.B interpreted results; K.E.C.B and M.N prepared figures; K.E.C.B., M.N., H.H and T.B writing - original draft; K.E.C.B., M.N., H.H., T.B., M.S., B.N.M., D.A.K., C.A.B., S.D.P and J.K.B Writing - review and editing. All authors have read and agreed to the final version of the manuscript. All authors have access to the raw data and B.N.M., C.A.B., S.D.P and J.K.B can verify the raw data in this study

## 8 Acknowledgments

The authors thank M.R. Jonker, L. Apperloo and G.J. Teitsma for their technical assistance in this study.

## 9 Data Availability Statement

Data available on request from the authors

