## Supplementary Data for "Substrate stiffness engineered to replicate disease conditions influence senescence and fibrotic responses in primary lung fibroblasts"

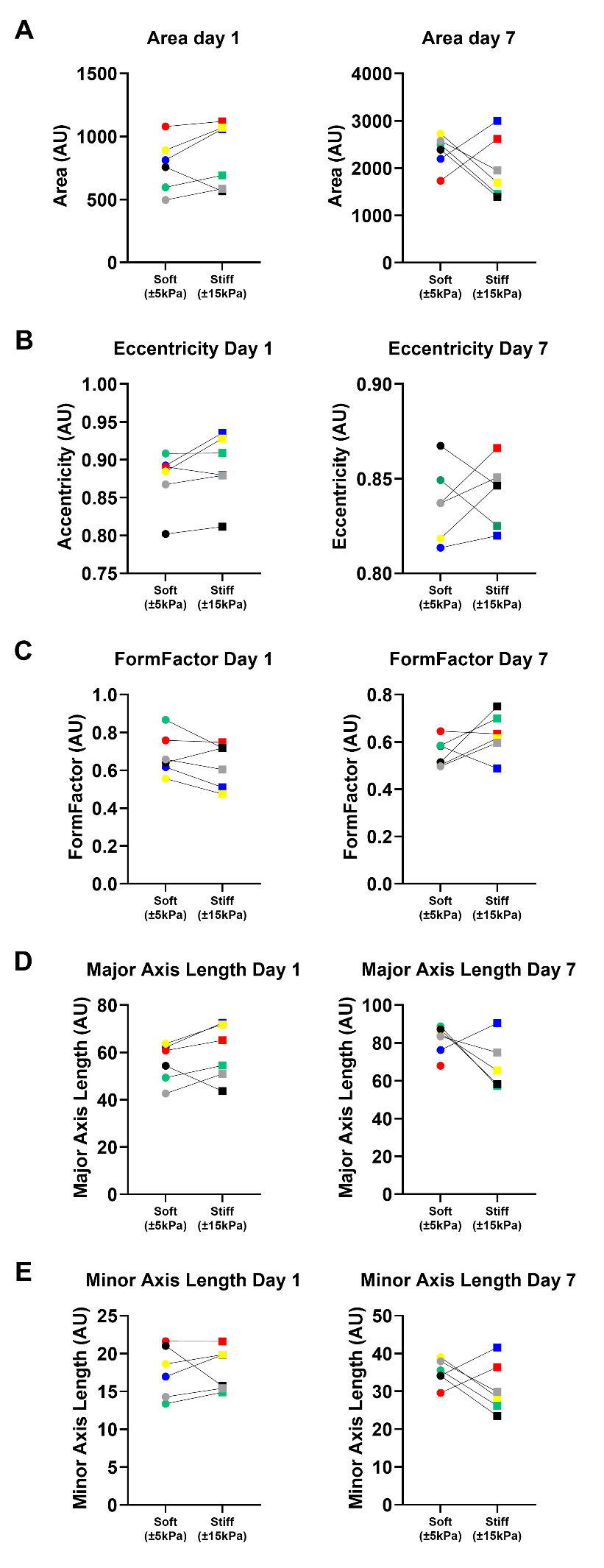

Supplementary figure 1; Cellular size and shape analysis of fibroblasts at day 1 and 7 on soft and stiff GelMA hydrogels. Quantification of cellular area (A), eccentricity (B), Formfactor (C, how defined the outline of object is) and Major and minor axis length (D-Ehow o) (n = 6). Each colour is an unique fibroblast donor (Blue, Yellow, Red, Yellow, Green, Black and Grey)
